# Large scale function-based genome prospecting for industrial traits applied to 1,3-propanediol production

**DOI:** 10.1101/2021.08.25.457110

**Authors:** Jasper J. Koehorst, Nikolaos Strepis, Sanne de Graaf, Alfons J. M. Stams, Diana Z. Sousa, Peter J. Schaap

**Affiliations:** Laboratory of Systems and Synthetic Biology | Department of Agrotechnology and Food Sciences, Wageningen University and Research, Wageningen, the Netherlands; Laboratory of Microbiology | Department of Agrotechnology and Food Sciences, Wageningen University and Research, Wageningen, the Netherlands; Centre of Biological Engineering, University of Minho, 4710-057 Braga, Portugal

**Author notes:** Equal contribution (Jasper J. Koehorst, Nikolaos Strepis).

**Keywords:** genome prospecting, trait, big data, Semantic Web, Knowledge and Discovery in Databases, 1,3-propanediol

## Abstract

Due to the success of next-generation sequencing, there has been a vast build-up of microbial genomes in the public repositories. FAIR genome prospecting of this huge genomic potential for biotechnological benefiting, require new efficient and flexible methods. In this study, Semantic Web technologies are applied to develop a function-based genome mining approach that follows a knowledge and discovery in database (KDD) protocol. Focusing on the industrial important trait of 1,3-propanediol (1,3-PD) production 187 new candidate species were identified. Furthermore, the genetic architecture of the particular trait was resolved, and persistent domains identified.

## Introduction

Next generation sequencing (NGS) technologies have turned the publicly available genome repositories into data-rich scientific resources that currently contain structural and functional genome information of hundred of thousands bacterial genomes (1) available for further exploitation for medical, environmental and biotechnological purposes. While the biotechnology field has embraced the use of omics technologies in biotech innovation in multiple ways, functional screening of genomic resources for industrially relevant traits at a large scale remains challenging. NGS data generation has caused manual curation to be outpaced rapidly and currently more than 99% of the functional predictions in UniProt are based on automated computational predictions (2). The accuracy of these computationally inferred functional genome annotations is largely unknown due to inconsistent naming of protein functions, use of annotation pipelines of various quality, over-prediction due to the use of generic, nonstandardised, annotation acceptance thresholds (3), and lack of data provenance in general. This results in a degree of interoperability lower than required for efficient function based genome prospecting (4). As the predicted gene and protein sequences, underlying the functional annotations are available in the standardised FASTA format, ‘bottom-up’ approaches that start with a gene or protein sequence in an effort to find a corresponding function in genome(s) of interest are normally used. However, over larger phylogenetic distances, strategies that use sequence-based clustering algorithms are hampered by accumulation of evolutionary signals, lateral gene transfer, gene fusion/fission events and domain expansions (5). Furthermore, when applied at larger scales, such strategies suffer from high computational time and memory requirements that scale quadratically with the number of genome sequences to be compared (6). A robust function-based genome screening requires a sufficiently high degree of interoperability meaning that functional information can be directly compared on the basis of a pre-established syntactic interoperable genome annotation and that any computational functional prediction is directly linked to its provenance. To accomplish this, we have developed SAPP, a semantic annotation infrastructure supporting FAIR computational genomics (7). SAPP uses the Genome Biology Ontology Language (GBOL) as syntax (8) and automatically predicts, tracks and stores structural and functional genome predictions and associated data- and element-wise provenance in RDF, the W3C standard for representing linked data (9). SAPP uses protein domain architectures to systematically describe protein functions (10). Demonstrating a high level of scalability, this set up has already been successfully used in an integrated analysis of the functional landscape of 432 Pseudomonas strains (4). In this study, interoperable genome annotations are used in a systematic function-based in silico screening for bacterial species that potentially can convert the bio-refinery by-product glycerol in 1,3-propanediol (1,3-PD) (11, 12). 1,3-PD is an important precursor of biomaterials and it is currently used as a monomer for novel polyester and biodegradable plastics, such as polytrimethylene terephthalate (11). 1,3-PD is a typical product of anaerobic glycerol fermentation and only a limited number of species, mostly enterobacteria, are known to form it. The first step, dehydration of glycerol to 3-hydroxypropionaldehyde (3-HPA) is mediated by a vitamin B12-dependent glycerol dehydratase although an oxygen sensitive, B12-independent alternative enzyme has also been reported (13). The second step reduces the toxic 3-HPA intermediate to 1,3-PD using the (NADH)+H+ dependent 1,3-propanediol-oxydoreductase (PDOR) regenerating NAD+ (12).

For large scale genome prospecting a collection of 84,308 publicly available bacterial genomes were loaded in the SAPP semantic framework, systematically structurally and functionally annotated and subsequently mined for candidate 1,3-PD producers using a knowledge discovery in databases approach (KDD) (14). Overall, the systematic analysis increased our knowledge on the genetic architecture of this metabolic trait, in terms of the overall domain composition, domain order, distribution, persistence and essentiality. The approach suggested that, in addition to the some 30 producers reported in literature (table 1), at least 187 other genome sequenced species potentially have this trait.

**Table 1.** Overview of known 1,3-PD producers.

| Organism | Genome assembly ID | B12<br>(in silico prediction) | DAK configuration | 1,3-PD<br>(mol/mol) | Acetate<br>(mol/mol) | Reference |
| --- | --- | --- | --- | --- | --- | --- |
| *Clostridium perfringens* str. 13 | GCA_000009685.1 | Dependent | DAK1/DAK1_2 |  |  |  |
| *Clostridium butyricum* E4 | GCA_000182605.1 | Independent | DAK1/ DAK1_2 | 0.58 | 0.03 | Petitdémange et al., 1995 |
| *Klebsiella michiganensis* strain M5al ==oxytoca | GCA_000308735.2 | (in-) dependent | DAK1 | 0.41 | 0.03 | Yang et al., 2007 |
| *Citrobacter freundii* ATCC 8090 | GCA_000312465.1 | Dependent | DAK1 | 0.65 | 0.22 | Barbirato et al., 1998 |
| *Citrobacter freundii* ATCC 8090 | GCA_000734905.1 | Dependent | DAK1 | 0.65 | 0.22 | Barbirato et al., 1998 |
| *Clostridium pasteurianum* ATCC 6013= DSM 525 | GCA_000330945.1 | Dependent | DAK1/ DAK1_2 | 0.5 | 0.12 | Taconi et al., 2009 |
| *Halanaerobium saccharolyticum* DSM 6643 | GCA_000350165.1 | Dependent | DAK1/ DAK1_2 | 0.41 | 0.26 | Kivistö et al., 2012 |
| *Clostridium diolis* DSM 15410 | GCA_000409695.1 | Independent | DAK1/ DAK1_2 | 0.58 | 0.12 | Otte et al., 2009 |
| *Klebsiella pneumoniae* ATCC 25955 | GCA_000409715.1 | Dependent | DAK1/ | 0.65 | 0.22 | Barbirato et al., 1998 |
| *Clostridium butyricum* VPI3266 = DSM 10702 | GCA_000409755.1 | Independent | DAK1/ DAK1_2 | 0.6 | 0.01 | Gonzalez-Pajuelo et al., 2004 |
| *Raoultella planticola* DR3 = DSM ATCC 33531 | GCA_000735435.1 | Independent (no alc) | DAK1 | N.D. | 0.15 | Jarvis et al., 1997 |
| *Klebsiella pneumoniae* DSM 2026 = ATCC 15380 | GCA_000949515.1 | Dependent | DAK1 | 0.62 | 0.09 | Mu et al., 2006 |
| *Clostridium beijerinckii* NRRL B-593 | GCA_002006325.1 | Independent (no alc) | DAK1_2 | 0.59 | 0 | Gungormusler et al., 2011 |
| *Trichococcus* strain ES5 | GCA_900067165.1[NS1] |  | DAK1/ DAK1_2 | 0.66 |  | Streps 2020 |
| *Caloramator viterbensis* DSM 13723T |  |  |  | 0.69 | 0.21 | Seyfried et al 2002 |
| *Clostridium butyricum* AKR102a |  |  |  | 0.52 | N.D. | Wilkens et al., 2012 |
| *Clostridium butyricum* CNCM 1211 |  |  |  | 0.64 | 0.11 | Barbirato et al., 1998 |
| *Clostridium butyricum* DSM 5431 |  |  |  | 0.51 |  | Rehman et al., 2008 |
| *Clostridium butyricum* E5 |  |  |  | 0.54 | 0.08 | Petitdémange et al., 1995 |
| *Clostridium butyricum* F2b |  |  |  | 0.63 | 0.13 | Papanikolaou et al., 2004 |
| *Clostridium butyricum* VPI 1718 |  |  |  | 0.68 | 0.04 | Chatzifragkou et al., 2011 |
| *Clostridium multifementans* DSM 5431 |  |  |  | 0.4 | 0.01 | Biebl et al., 1992 |
| *Klebsiella pneumoniae* DSM4799 |  |  |  | 0.45 | N.D. | Jun et al., 2010 |
| *Klebsiella pneumoniae* HR526 |  |  |  | 0.4 | N.D. | Xu et al., 2009 |
| *Klebsiella pneumoniae* XJ-Li |  |  |  | 0.61 | 0.05 | Zhan et al., 2007 |
| *Lactobacillus brevis* N1E9.3.3 |  |  |  | 0.46 | 0.07 | Vivek et al., 2016 |
| *Lactobacillus buchneri* B 190 |  |  |  | 0.27 | 0.18 | Schutz et al., 1984 |
| *Lactobacillus panis* PM1 |  |  |  |  |  | Tae Sun Kang (2014) |
| *Pantoea agglomerans* CNCM-1210 |  |  |  | 0.61 | 0.16 | Barbirato et al., 1998 |
| *Shimwellia blattae* ATCC 33430 |  |  |  | 0.45 g/g |  | Rodríguez 2016 |
| *Clostridium beijerinckii* DSM 791 |  |  |  | 0.55 (gg 1) |  | Pradima 2017 |

## Results

### Development of the genome mining workflow

To cope with the influx of biological input data, we followed the Knowledge Discovery in Databases multi step process model (14), which was adjusted to the specific needs of being able to prospect genome data at a large scale. The KDD process includes multiple steps with the first step being to capture relevant prior knowledge. In the next step, a selection of data is used as a training set to develop a screening procedure followed by data mining at a large scale and finally the evaluation and interpretation of the results.

### Incorporation of prior knowledge

In 1,3-PD production, two pathways, an oxidative and reductive pathway, are used in parallel for dissimilation of glycerol. In the oxidative pathway, glycerol is first dehydrogenated to dihydroxyacetone by a NAD+-linked glycerol dehydrogenase, and then converted to dihydroxyacetone phosphate by an ATP-dependent dihydroxyacetone kinase (15). Additionally, two alternative reductive pathways, one dependent on vitamin B12, the other not, are used for regenerating NAD+ with 1.3-PD as byproduct. For validation of the search strategy and to gain further insights into the genetics and the domains involved in 1,3-PD formation a list of known 1,3 PD producing strains was obtained for cross-validation. From literature some 30 strains have been reported to produce 1,3-PD (Table 1). For training purposes genome sequences from 13 different strains could be obtained from the EBI-ENA data warehouse (1) as not for all of them genome sequences were available. For *Citrobac-terfreundii* ATCC8090 two draft genome sequences of comparable quality were available and initially both were added to the training set increasing the total number of genomes.

### Data transformation

To obtain a high degree of interoperability the selected genome sequences were de novo structurally and functionally annotated using the Semantic annotation Platform with Provenance (SAPP) (7) using Prodigal (16) for gene prediction and InterProScan (17) for protein annotation. Protein domain architectures were used to systematically describe the encoded protein functions (10). Structural and functional genome annotations were subsequently transformed into Linked Data fragments using the Resource Description Framework (RDF) as a data-metadata model (7, 18).

### A function-based search strategy for the oxidative branch

The first two enzymes essential for the oxidative pathway involved in the production of 1,3 PD are glycerol dehydrogenase which is dependent on NADH and produces dihydroxyacetone (DHA) and DHA-kinase, which subsequently phosphorylates DHA. These key reactions are represented by four Pfam domains; Fe-ADH (PF00465) for glycerol dehydrogenase and DAK1, (PF02733) or DAK1_2, PF13684) both capturing the kinase domain of the dihydroxyacetone kinase family in combination with DAK2, (PF02734), capturing phosphatase domain of the dihydroxyacetone kinase family (Figure 1). A pattern recognition analysis was performed (see methods section for details) for DAK2 domain neighbours in the training set database followed by a Boolean DAK1/DAK2 or DAK1_2/DAK2 proximity search in the search results. Pattern recognition showed that the 14 genomes of the training data set encoded at least one dihydroxyacetone kinase family protein with either a DAK1/DAK2 or DAK1_2/DAK2 protein architecture; 13 strains coded for DAK1/DAK2, 8 strains for DAK1_2/DAK2 and 7 strains encoded for both configurations (Table 1). In addition, other domains were identified that clustered together with the DAK configuration such as an Alkaline shock protein, a phosphotransferase system component and an alcohol dehydrogenase domain. Another copy of this domain is an important factor for the 1,3 PD pathway (Table 2).

**Fig. 1.**
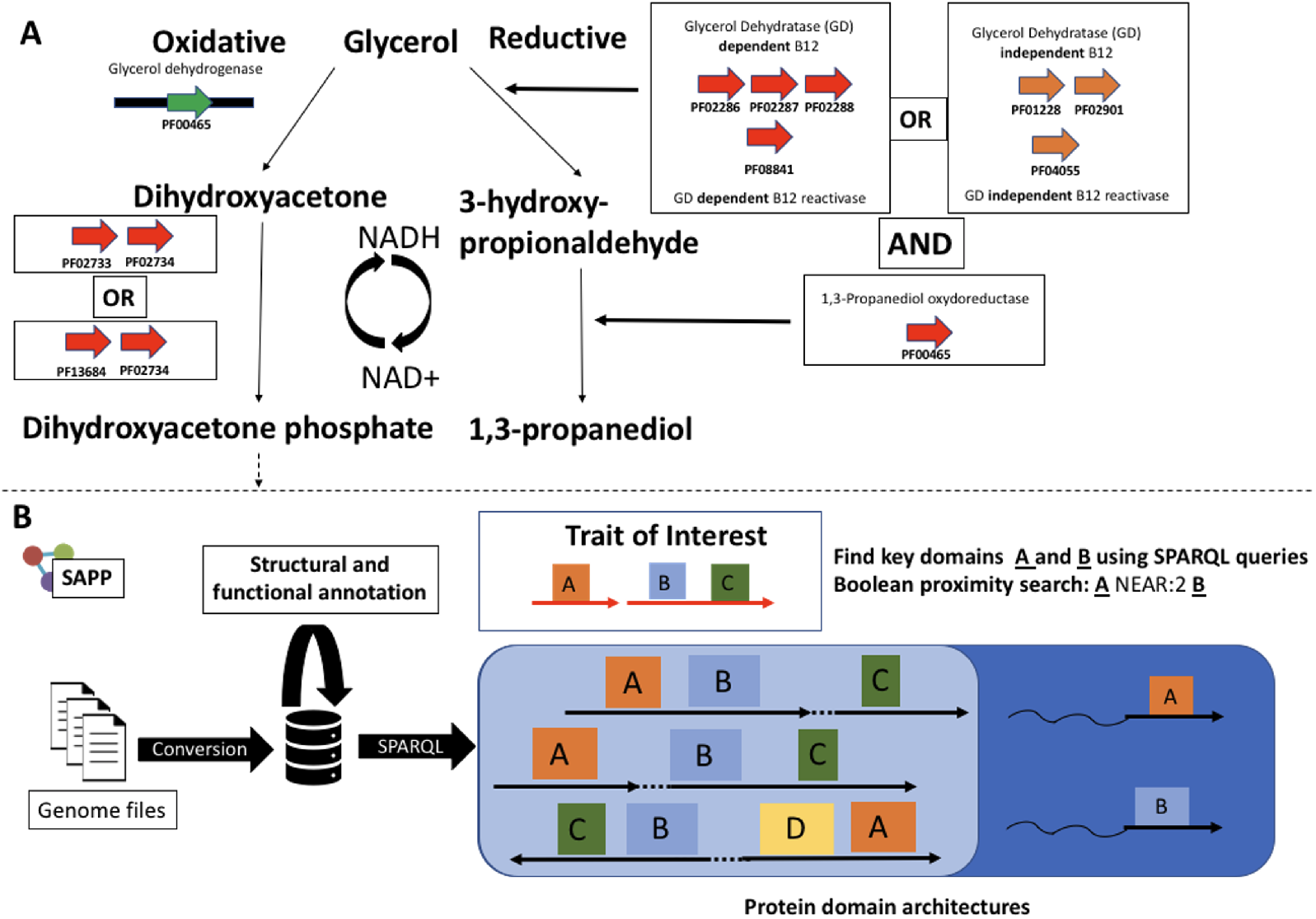
Search strategy for 1,3-Propanediol candidate species: **(A)** Key domains in the main pathway for B12 dependent bio-conversion of glycerol to 1,3-propanediol are indicated in red. Key domains for the alternative B12-independent reductive pathway are indicated in orange. Note the generic iron alcohol domain is present in both the oxidative and reductive branch. **(B)** Generalised functional based search strategy for traits using SAPP. Genome sequences in standard format are converted and complemented with structural and functional annotation. Pattern recognition strategies are deployed to identify domains of interest (dark blue) and complemented with proximity searches (light blue) to find key domain enriched regions.

**Table 2.** Reference set DAK analysis is this needed, or should we modify the text to make it fit?

| Domain | PatternFrequency | support | JaccardIndex | Domain description |
| --- | --- | --- | --- | --- |
| PF02734 | 32 | 1 | 1 | DhaL domain |
| PF02733 | 19 | 0.59 | 0.93 | DhaK domain |
| PF13684 | 8 | 0.25 | 0.75 | Uncharacterised protein, DhaK domain |
| PF03780 | 8 | 0.25 | 0.41 | Alkaline shock protein Asp23 |
| PF03610 | 16 | 0.5 | 0.11 | Phosphotransferase system, mannose-type IIA component |
| PF00465 | 8 | 0.25 | 0.05 | Alcohol dehydrogenase, iron-type |

### Development of a function-based search strategy for the reductive branch

For NAD+ regeneration and 1,3-PD production two alternative pathway are known, a well-studied B12-dependent pathway and a lesser studied oxygen sensitive B12-independent pathway (figure 1). Pattern recognition analysis that was used to obtain the DAK configuration was used to obtain the domain persistency and copy number of key domain classes in the two alternative pathways among the genomes of the training set. The iron containing aldehyde reductase/alcohol dehydrogenase domain (PF00465), is present in both the oxidative and reductive branch but functions in the reductive branch as an aldehyde reductase. The three dehydratase Pfam domains signifying the B12-dependent pathway, PF02287 (small subunit), PF02288 (medium subunit), and PF02286 (large subunit) have a domain class persistency of 0,6 with, when present, on average per genome 1,7 (large and small subunit) or 3,7 copies (medium subunit) in the training set. The diol dehydratase reactivase ATPase-like domain PF08841 has a persistency of 0,6 and a copy number of 1. Note that only four strains in the training data set were molecularly characterized. The degree of persistency of these key domains of the B12-dependent pathway in the training set therefore suggests that multiple 1,3-PD-producing strains of the training set may actually not use the B12-dependent pathway for 1,3 PD-production. Alternatively, a B12-independent glycerol dehydratase function is reported to be involved in 1,3-PD production (13) and consists of a glycine radical (PF01228) and a pyruvate formate lyase-like domain (PF02901). Both domains are present in all 13 genomes of the training data set with a relative high copy number of 6. The persistency of the radical SAM superfamily domain, a key part of the B12-independent glycerol dehydratase reactivase enzyme (19), was also 1 with an average copy number of 32 indicating that these three domains are ubiquitous and promiscuous and normally also used to support other functionalities (Table 3). As the high copy number of persistent signifying domains of both the B12-dependent and B12-independent reductive branch suggests that they play roles in multiple processes, a Boolean multi-compound proximity search was developed for the reductive branch. In this approach, the physical co-localization of the persistent signifying domain classes in the respective genomes is included in the search. From the list of natural producers with a molecularly characterized 1,3-PD operon, the size of 1,3-PD operon was estimated to encompass approximately 18,000 bases. Furthermore, signifying domains could be found on both strands due to the use of internal promoters.

**Table 3.** B12-independent

| Domain | PatternFrequency | Support | JaccardIndex | Domain description |
| --- | --- | --- | --- | --- |
| PF00465 | 4 | 1 | 0.03 | Alcohol dehydrogenase, iron-type |
| PF01228 | 4 | 1 | 0.07 | Glycine radical domain |
| PF02901 | 4 | 1 | 0.11 | Pyruvate formate lyase domain |
| PF04055 | 4 | 1 | 0.01 | Radical SAM |
| PF00465 | 4 | 1 | 0.03 | Alcohol dehydrogenase, iron-type |
| PF13353 | 3 | 0.75 | 0.03 | 4Fe-4S single cluster domain |
| PF03319 | 1 | 0.25 | 0.08 | Ethanolamine utilization protein EutN/carboxysome structural protein Ccml |
| PF06130 | 1 | 0.25 | 0.12 | Phosphate propanoyltransferase |

For the B12-independent pathway search criteria were based on prior knowledge obtained from a single characterized strain (13). Here four domains were specified (PF00465, PF01228, PF02901, PF04055) extending 20.000 base pairs up and downstream of the alternative dehydratase domains. Furthermore, taking internal promoters into account, domains were allowed to be present on both strands when genes are facing in the opposite direction in case of bidirectional promoter.

The B12-dependent pathway should contain at least five signature domains (PF00465, PF02286, PF02287, PF02288, PF08841) and were analyzed with the same restrictions as applied to the independent pathway.

Including the signifying domains used in the query, the core of the B12-dependent 1,3-PD operon may consist of eight domains. Other highly persistent domains (Support level > 0.5) enriched within the syntenic region are ‘Cobalamin adenosyltransferase’, (PF01923), ‘Haem-degrading’ (PF03928), ‘Alcohol dehydrogenase, iron-type’ (PF00465), and the ‘Microcompartment protein’ (PF00936), (see Table 4).

**Table 4.** Frequent pattern analysis on reference genomes for B12-dependent 1,3 PD and a minimum support value of 0.5

| Domain | PatternFrequency | Support | JaccardIndex | Domain description |
| --- | --- | --- | --- | --- |
| PF02288 | 12 | 1 | 0.86 | Diol/glycerol dehydratase/dehydratase reactivating factor |
| PF08841 | 12 | 1 | 1 | Diol dehydratase reactivase ATPase-like domain |
| PF02286 | 12 | 1 | 1 | Diol/glycerol dehydratase, large subunit |
| PF02287 | 12 | 1 | 1 | Propanediol/glycerol dehydratase, small subunit |
| PF01923 | 8 | 0.67 | 0.4 | Adenosylcobalamin biosynthesis, ATP:cob(I)alamin adenosyltransferase-like |
| PF03928 | 8 | 0.67 | 0.59 | Haem-degrading |
| PF00465 | 8 | 0.67 | 0.08 | Alcohol dehydrogenase, iron-type |
| PF00936 | 6 | 0.5 | 0.35 | Microcompartment protein, bacteria |

### Data preparation

To be able to mine a large set of publicly available genomes for the presence of the vitamin B12 variant 1,3-PD operon, 84,308 bacterial genomes containing 51 phyla, 64 classes, 145 orders, 335 families, 1,126 genera and 2,661 species were downloaded from the EBI-ENA data warehouse. The SAPP annotation platform was subsequently used for *de novo* structural and functional annotation of these genomes resulting in a semantic database of 300,196,266 predicted protein encoding genes linked to the corresponding protein sequences, predicted protein domain architectures and provenance.

### Data Mining for candidate 1,3 PD producers

Two parallel pathways must be present to convert glycerol to 1,3-PD. Data mining for candidate 1,3-PD producers was therefore split in two steps. First, the search space was reduced by searching for the presence of a possible oxidative branch. In the second step, the reduced search space was searched for the presence of a possible reductive branch. As the molecular knowledge obtained for B12-independent pathway is based on the characterization of a single *Clostridium butyricum* strain, data mining for the reductive branch focused on finding candidate B12-dependent 1,3 PD producers.

#### Oxidative branch

Following the function-based search strategy outlined above and summarised in figure 1, a proximity search with the two DAK configurations in parallel resulted in a 41% reduction of the search space. The thus reduced search space contained 49,110 genomes assigned to 3,526 species from 976 genera. The most abundant species were *Listeria monocytogenes* (5521), *Streptococcus pneumoniae* (6305), *Escherichia coli* (7577) and *Staphylococcus aureus* (10232). The reduced search space was used as an input for a proximity search for the presence of the B12-dependent reductive pathway.

#### Reductive branch

Identification of B12-dependent strains according to the criteria obtained from the training set and based upon the results obtained from the oxidative branch, resulted in a list of 247 species (5,538 strains). Strains of *Klebsiella pneumoniae* (1,637) and *Listeria monocytogenes* (1,788), both pathogenic species were over-represented as well as *Escherichia coli* (1,067). An overview of identified species can be found in Supplementary table S1. The most frequently observed additional syntenic domains showing a persistency of 0.75 and above are shown in Table 6.

**Table 5.** Oxidative on genome resource

| Domain | PatternFrequency | support | JaccardIndex | Domain description |
| --- | --- | --- | --- | --- |
| PF02734 | 74929 | 1 | 1 | DhaL domain |
| PF02733 | 38482 | 0.51 | 0.91 | DhaK domain |
| PF13684 | 32142 | 0.43 | 0.97 | Uncharacterised protein, DhaK domain |
| PF03610 | 27381 | 0.37 | 0.15 | Phosphotransferase system, mannose-type IIA component |
| PF03780 | 23828 | 0.32 | 0.34 | Alkaline shock protein Asp23 |

**Table 6.**
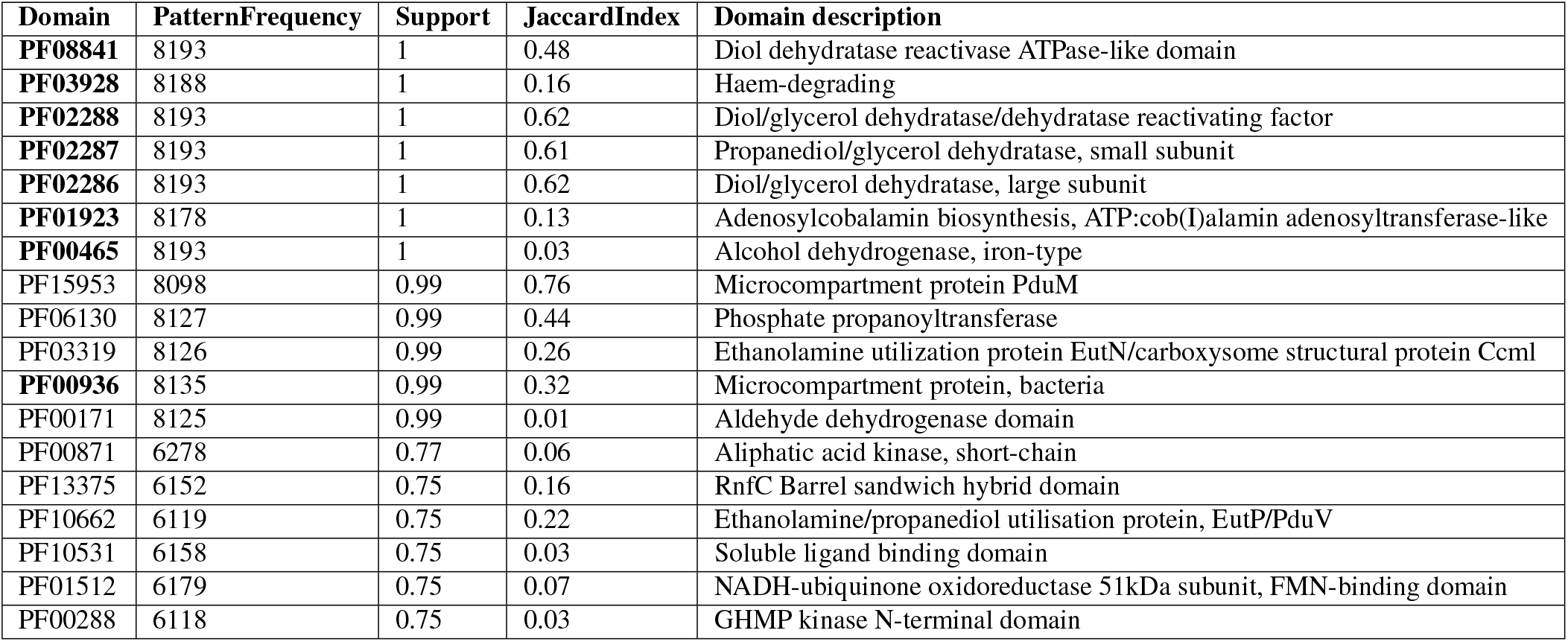
Pattern frequency analysis for the reductive branch on genomes obtained from the oxidative analysis. Protein domains in **bold** are reoccurring from the training set.

For further analysis, strains were collapsed in species groups and each species group was treated as one. At species level, domain connectivity (figure 2) and persistency of the B12-dependent 1,3-PD trait showed a high degree of similarity with the training set (Table 6).

**Fig. 2.**
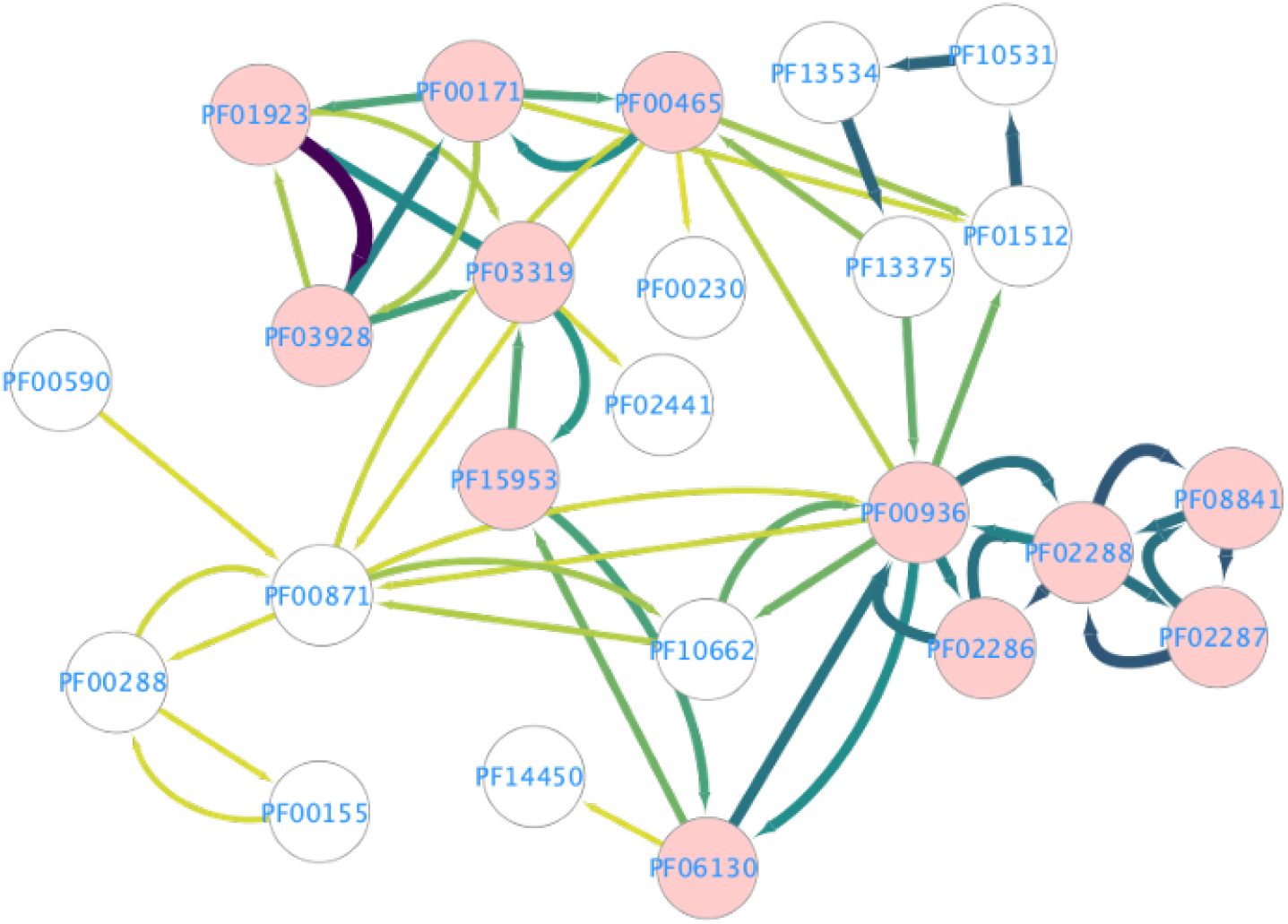
Protein domain connectivity order. Arrows indicate the order in which protein domains were identified in operonic structures. The darker and thicker the arrow indicates that this connection was more observed.

### Additional constraints

As the candidate species are depending on vitamin B12 for the production of 1,3-PD it could be beneficial to additionally select for the ability to synthesize vitamin B12. Secondly, the best known natural B12 dependent 1,3-PD producers are considered to be pathogenic or opportunistic pathogenic microorganisms such as *Klebsiella pneumonia* (20).

Following the strategy outlined above a function-based screening was developed for the distribution of the trait for vitamin B12 synthesis among the candidate species. Using a multi-domain approach containing all domain signatures reuired for vitamine B12 sysnthesis a vitamin B12 (21) biosynthesis pathway was detected in 86 of the 187 candidate species using a multi-domain approach containing all domain signatures required for vitamin B12 synthesis. In order to obtain the bio-safety level of each candidate species, the BacDive database was queried (22). Forty-seven candidate species were classified at bio-safety level 2 and higher and 48 species were considered to be non-pathogenic (biosafety level 1) while 92 species remained unclassified. The bio-safety level 1 label was combined with the ability for vitamin B12 synthesis with resulted in the following candidate species; *Anaeromusa acidaminophila, Anaerosalibacter massiliensis, Blautia obeum, Clostridium drakei, Clostridium magnum, Geobacillus thermoglucosidasius, Intestinimonas butyriciproducens* and *Thermincola ferriacetica*, (see Table suppl S1).

## Discussion

Bioprospecting entails the systematic search for economically valuable genetic and biochemical resources from nature (23). For genome prospecting, the *in silico* mining of sequenced genomes and metagenomes for new biotechnologically relevant proteins and enzymes is a continuously evolving field. Two approaches can be applied, a “top-down” approach that begin by searching for a function and is followed by identification of the corresponding gene(s) and a “bottom-up” approach that start with a gene of interest in an effort to find a corresponding function in genome(s) of interest. For bottom-up approaches many sequence similarity tools such as Blast (24) exist. Bottom-up studies usually start with a selection of (meta)genome sequences followed by sequence similarity-based clustering and selection of candidate sequences and re-annotation of interesting candidates thereby avoiding ambiguity related problems in current functional annotations. For bacterial species however, *a priori*, gene fusion-fission events can be expected (25) hampering sequence similarity-based detection and clustering of multidomain proteins encoding genes, while the same domains may also be present in multiple proteins. When searching for polygenic traits these problems are aggravated and thus a function-based approach searching for key functions may be more effective especially when there is insufficient understanding of the genetic architecture of trait in terms of the minimal number of genes required for the trait and function of the domains encoded by the different genes. In order to overcome ambiguity related problems in functional annotations, SAPP identifies and annotates protein function based on domain architecture. Protein domains have been shown to provide an accurate representation of the functional capabilities of a protein (10). To overcome ambiguity related problems in functional annotations SAPP identifies and annotates protein function based on domain architecture. As profile hidden Markov Models (HMM), which favour in their scoring functionally important sites, are used for the identification of protein domains, statistically robust annotation profiles can be obtained over large phylogenetic distances. KDD is a multi-step process. Crucia steps are data standardisation and feature extraction for pattern searching. By using protein domain architectures as proxy for protein functions, a high degree of standardisation is obtained. As protein domain architectures can be directly transformed in highly interoperable strings of Pfam domain identifiers, a “top-down” functional screening can be done efficiently.

Applying the KDD approach on Linked Data allows for validation of initial results in multiple ways and for iteration after modification. Driven by a strong need to be able to integrate and analyse biological datasets across databases, there has been a considerable increase in the adoption of Semantic Web technologies in the life-sciences (26). However, a SPARQL endpoint for phenotype data is only available via WikiData (27). Other resources for phenotype data such as BacDive, here used to obtain bio-safety levels of candidate species, do provide API’s to mine their data but currently cannot be directly queried using SPARQL. With the growing importance of Semantic Web technologies for the life sciences interoperability levels on all aspect will increase, enhancing mining possibilities, aiding the discovery of new traits in an unprecedented pace.

By using molecular knowledge obtained from molecular characterisations of the 1,3-PD operon of four species and validating and iterating the search pattern on a training data set of genome sequences of twelve known 1,3-PD producers, a large collection of 84,308 publicly available bacterial could be efficiently mined in a top-down approach yielding 187 new candidate species some of which could be confirmed by literature (28–32).

## Conclusions

Through transformation of genomic data into a FAIR linked-data format, iterative function-based approaches can be developed to mine the large genome repositories. By presenting functional annotation as unambiguous protein domain architectures a high level of interoperability is obtained allowing for the development of efficient function-based top-down searches not limited to supervised trait identification.

## Materials and methods

### Data annotation and mining

Bacterial genomes were downloaded from the ENA database using the enaBrowser-Tools (1). All downloaded microorganisms were converted into a semantic repository using the annotation platform based on functional analysis (SAPP) (7). All genomes were *de novo* structurally re-annotated using Prodigal (16) and functionally annotated using Pfam from InterProScan (17). Each genome was stored as an individual graph database using the HDT library and were processed using a domain pattern recognition software that was developed to aid in the data processing (33, 34). This application constructs operon-like structures around one or more protein domains from genomic data and discovers patterns in the present protein domains using a Frequent Pattern Mining and a Latent Dirichlet Allocation approach.

### Pattern recognition

The application that was used for pattern recognition has been developed in Java 8 and uses Scala version 2.11. The annotated genomes are provided as RDF data in HDT format allowing to easily scale beyond the size limitations of current graph databases. Spark Core has been implemented to enable parallelization for the processing of the genomes. The Frequent Pattern Mining and Latent Dirichlet allocation algorithms from the Apache SPARK MLlib Machine Learning Library, are applied to analyse patterns in the domain architecture of operons Meng2016. The support value of a pattern with one domain is a measure for the local persistency of that specific domain. The support value is defined as the number of regions where the domain occurs divided by the total number of regions.

### Oxidative branch identification

To identify strains containing the oxidative branch as shown in figure 1, DAK domains were processed using the domain pattern recognition software. Genes containing DAK1 (PF02733) or DAK1_2 (PF13684) in combination with DAK2, (PF02734), 95% gene confidence according to prodigal and neighbour linking was applied to identify strains capable for growth on glycerol.

### Reductive branch identification

Strains that contained the oxidative branch were further investigated for the reductive branch using domains according to the B12 dependent or independent pathway. Region identification was performed using PF02287 and 20,000 base pairs up and downstream. Regions containing all domains of interest were further investigated for domain occurrence and synteny.

### Reductive branch identification

Strains that contained the oxidative branch were further investigated for the reductive branch using domains according to the B12 dependent or independent pathway. Region identification was performed using PF02287 and 20,000 base pairs up and downstream. These regions were modified to only retain genes that form an operon-like structure. The biggest intergenic region allowed between two genes on the same strand was 50 nucleotides. To also include the possibility of a bidirectional promoter, a gap of 500 (or less) nucleotides is allowed between to stretches of genes on the reverse and forward strand. To prevent identical regions created around a gene which contains multiple copies of the PF02287 or different genes containing PF02287, regions that (partly) overlap are merged into one. The operon-like structure was preserved.

### B12 synthesis identification

Phylogenetic information was complemented with phenotype information through Bac-Dive (22). The BacDive resource was parsed and each entry record was transformed into RDF allowing integration of genotype and phenotype information. SPARQL queries were used to retrieve information with regards to pathogenicity and temperature.

## ACKNOWLEDGEMENTS

This research was supported by the European Research Council under the European Union’s Seventh Framework Programme (FP/2007-2013) / ERC Grant Agreement (323009) and by the Gravitation grant (024.002.002) of the Netherlands Ministry of Education, Culture and Science. The work conducted by the U.S. Department of Energy Joint Genome Institute (DOE-JGI), a DOE Office of Science User Facility, was supported by the Office of Science of the DOE under Contract No. DE-AC02-05CH11231.IBISBA 1.0 - Project ID: 730976 - Funded under: H2020-EU.1.4.1.2. - Integrating and opening existing national and regional research infrastructures of European interest. This work was carried out on the Dutch national e-infrastructure with the support of the SURF foundation.

## Supplementary

**Table S1.** Overview of species identified to have the potential of producing 1,3 PD

|  |  |  |
| --- | --- | --- |
| Anaeromusa acidaminophila | Eubacteriaceae bacterium CHKCI004 | Lactobacillus kimchicus |
| Anaerosalibacter massiliensis | Eubacterium sp. 14-2 | Lactobacillus kisonensis |
| Aneurinibacillus migulanus | Eubacterium sp. An11 | Lactobacillus mellifer |
| Aneurinibacillus sp. UBA3580 | Eubacterium sp. An3 | Lactobacillus namurensis |
| Bacillaceae bacterium MTCC 10057 | Eubacterium sp. UBA3279 | Lactobacillus panis |
| Bacillus azotoformans | Eubacterium sp. UBA3326 | Lactobacillus paracollinoides |
| Bacillus massiliosenegalensis | Eubacterium sp. UBA7134 | Lactobacillus pobuzihii |
| Bacillus sp. 7504-2 | Firmicutes bacterium UBA3567 | Lactobacillus rapi |
| Bacteroides fragilis | Firmicutes bacterium UBA3570 | Lactobacillus rennini |
| Blautia obeum | Firmicutes bacterium UBA3573 | Lactobacillus reuteri |
| Blautia schinkii | Flavonifractor plautii | Lactobacillus rossiae |
| Blautia sp. UBA2945 | Flavonifractor sp. An306 | Lactobacillus silagei |
| Carnobacteriaceae bacterium UBA7837 | Flavonifractor sp. An82 | Lactobacillus siliginis |
| Carnobacterium alterfunditum | Fusobacteriaceae bacterium UBA2433 | Lactobacillus similis |
| Carnobacterium funditum | Fusobacterium hwasookii | Lactobacillus spicheri |
| Cetobacterium somerae | Fusobacterium nucleatum | Lactobacillus versmoldensis |
| Citrobacter amalonaticus | Fusobacterium sp. CM1 | Listeria innocua |
| Citrobacter freundii complex | Fusobacterium sp. CM22 | Listeria ivanovii |
| Citrobacter koseri | Fusobacterium sp. HMSC064B11 | Listeria monocytogenes |
| Citrobacter rodentium | Fusobacterium sp. HMSC073F01 | Listeria seeligeri |
| Citrobacter sp. 30_2 | Fusobacterium sp. OBRC1 | Listeria welshimeri |
| Citrobacter sp. A1 | Fusobacterium ulcerans | Metakosakonia massiliensis |
| Citrobacter sp. CFSAN044567 | Fusobacterium varium | Mycobacterium vaccae |
| Citrobacter sp. KTE151 | Geobacillus sp. (strain Y4.1MC1) | Paraclostridium benzoelyticum |
| Citrobacter sp. KTE30 | Geobacillus thermoglucosidasius | Paraclostridium bifermentans |
| Citrobacter sp. KTE32 | Geosporobacter ferrireducens | Pediococcus acidilactici |
| Citrobacter sp. L17 | Halanaerobium hydrogeniformans | Pediococcus clausenii |
| Clostridia bacterium UC5.1-2H11 | Halanaerobium saccharolyticum | Pediococcus pentosaceus |
| Clostridiaceae bacterium BRH_c20a | Ilyobacter polytropus | Pelosinus fermentans |
| Clostridiales bacterium DRI-13 | Intestinimonas butyriciproducens | Pelosinus sp. HCF1 |
| Clostridiales bacterium PH28_bin88 | Klebsiella aerogenes | Phytobacter ursingii |
| Clostridiales bacterium VE202-03 | Klebsiella michiganensis | Pluralibacter gergoviae |
| Clostridiales bacterium mt11 | Klebsiella oxytoca | Propionibacterium freudenreichii |
| Clostridium baratii | Klebsiella pneumoniae | Propionimicrobium sp. BV2F7 |
| Clostridium botulinum | Klebsiella quasipneumoniae | Pseudopropionibacterium propionicum |
| Clostridium drakei | Klebsiella sp. 10982 | Quasibacillus thermotolerans |
| Clostridium estertheticum | Klebsiella sp. 1_1_55 | Salmonella choleraesuis |
| Clostridium lundense | Klebsiella sp. AA405 | Sebaldella termitidis |
| Clostridium magnum | Klebsiella sp. AS10 | Serratia marcescens |
| Clostridium pasteurianum | Klebsiella sp. HMSC09D12 | Shigella boydii |
| Clostridium perfringens | Klebsiella sp. HMSC16A12 | Shigella flexneri |
| Enterobacter cloacae complex | Klebsiella sp. HMSC22F09 | Shigella sonnei |
| Enterobacter sp. 10-1 | Klebsiella sp. KGM-IMP216 | Sporomusa acidovorans |
| Enterobacter sp. Bisph2 | Klebsiella sp. KTE92 | Sporomusa sphaeroides |
| Enterobacter sp. GN02600 | Klebsiella sp. LTGPFAF-6F | Sporomusaceae bacterium UBA5961 |
| Enterobacteriaceae bacterium UBA2606 | Klebsiella sp. M5al | Sporomusaceae bacterium UBA5989 |
| Enterobacteriaceae bacterium UBA3163 | Klebsiella sp. OBRC7 | Sporomusaceae bacterium UBA7701 |
| Enterobacteriaceae bacterium UBA3516 | Klebsiella sp. PO552 | Sporomusaceae bacterium UBA864 |
| Enterobacteriaceae bacterium UBA4747 | Klebsiella sp. S1 | Streptococcus australis |
| Enterobacteriaceae bacterium UBA5935 | Klebsiella variicola | Streptococcus sanguinis |
| Enterobacteriaceae bacterium UBA869 | Klebsiella variicola CAG:634 | Streptococcus sp. AS14 |
| Enterobacteriaceae bacterium UBA898 | Kluyvera intermedia | Streptococcus sp. F0442 |
| Enterococcus avium | Lachnospiraceae bacterium 3-1 | Streptococcus suis |
| Enterococcus malodoratus | Lachnospiraceae bacterium 7_1_58FAA | Thermincola ferriacetica |
| Enterococcus mundtii | Lachnospiraceae bacterium A2 | Thermincola potens |
| Enterococcus pseudoavium | Lactobacillus brevis | Thermoanaerobacter sp. (strain X513) |
| Enterococcus raffinosus | Lactobacillus collinoides | Thermoanaerobacter sp. (strain X514) |
| Enterococcus sp. 3H8_DIV0648 | Lactobacillus coryniformis | Thermoanaerobacter sp. A7A |
| Enterococcus sp. HMSC066C04 | Lactobacillus curvatus | Thermoanaerobacterium thermosaccharolyticum |
| Enterococcus sp. HMSC072H05 | Lactobacillus diolivorans | Tissierellia bacterium S5-A11 |
| Enterococcus sp. HMSC29A04 | Lactobacillus fuchuensis | Eubacterium hallii |
| Escherichia coli | Lactobacillus ginsenosidimutans |  |
| Escherichia fergusonii | Lactobacillus graminis |  |

## Notes

### Competing Interest Statement

The authors have declared no competing interest.

